# N6-methyladenosine regulation of mRNA translation is essential for early human erythropoiesis

**DOI:** 10.1101/2025.11.10.687731

**Authors:** Daniel A. Kuppers, Sonali Arora, Cindy L. Wladyka, Ruiqi Ge, Shun Liu, Yong Peng, Rui Su, Anne Wilhite, Jianjun Chen, Chuan He, Andrew C. Hsieh, Patrick J. Paddison

**Author notes:** Correspondence: Daniel Kuppers or Patrick Paddison.

## Abstract

N6-methyladenosine (m^6^A) is an abundant modification of mRNA with important regulatory roles in normal and malignant hematopoiesis. We previously reported that in human erythroid leukemia (HEL) cells, m^6^A mRNA marking selectively regulates translation of essential erythropoiesis genes required for in vitro differentiation and human erythroid colony formation. Here, we further investigated the timing and nature of requirement for m^6^A-methyltransferase (MTase) activity during human erythropoiesis, using a standardized *in vitro* erythroid differentiation assay for hHSPCs. We identified two critical m^6^A regulated developmental windows in BFU-E and during the transition from CFU-E to proerythroblasts. These windows of m^6^A-MTase requirement coincide with rising global m^6^A levels, which peak in proerythroblasts. After proerythroblast formation, however, m^6^A -MTase activity is dispensable for differentiation, proliferation, and survival. In BFU-E, m^6^A-MTase promotes proliferation but is dispensable for differentiation, while, in CFU-E, both m^6^A -MTase and the YTHDF family of m^6^A readers are essential for differentiation to proerythroblasts. Mechanistically, in CFU-E, m^6^A MTase activity enhances translation of ribosomal and oxidative phosphorylation (OXPHOS) genes, thereby elevating global protein synthesis rates and enabling efficient erythroblast formation. We propose that this form of translational regulation by m^6^A emerged as an evolutionary adaptation to meet the high translational demands of human erythropoiesis.

## Introduction

Erythropoiesis is a highly orchestrated process that transforms hematopoietic stem and progenitor cells (HSPCs) into mature, enucleated red blood cells through successive stages of lineage commitment and terminal differentiation^1^. Each developmental transition is accompanied by precise coordination of transcriptional and translational programs that regulate proliferation, metabolism, and hemoglobinization. Disruption of these regulatory mechanisms results in ineffective erythropoiesis. Serumfree in vitro differentiation systems using human CD34⁺ HSPCs now permit detailed temporal analysis of erythroid development and enable dissection of stage-specific molecular requirements^2^.

Among post-transcriptional mechanisms that modulate gene expression, N^6^Methyladenosine (m^6^A) methylation has emerged as a major regulator of mRNA fate^3^. Deposited by the m^6^A methyltransferase (MTase) complex, with a core catalytic subunit of METTL3 and METTL14, and interpreted by various reader protein families, including YTHDF, YTHDC, and IGF2BP reader proteins, m^6^A regulates mRNA splicing, stability, and translation^3,4^. Recent studies have revealed that m^6^A is essential for maintaining hematopoietic stem cell self-renewal and lineage commitment in myelopoiesis and lymphopoiesis^5–10^. Our previous work demonstrated that in human erythroleukemia (HEL) cells, m^6^A selectively promotes translation of erythroid genes, including ribosomal proteins and chromatin regulators, thereby enabling erythroid differentiation and colony formation^11^. However, whether the same m^6^A-dependent translational mechanisms operate in primary human erythroid progenitors remains unclear.

Here, we define the requirement for m^6^A methyltransferase (m^6^A -MTase) activity during early human erythropoiesis using a standardized, serum-free CD34⁺ HSPC differentiation system^2^. We identify two critical developmental windows in which m^6^A MTase activity is essential: during BFU-E expansion and during the CFU-E–to–proerythroblast transition. Mechanistically, m^6^A methylation promotes translation of ribosomal and oxidative phosphorylation (OXPHOS) genes, thereby sustaining the biosynthetic and energetic demands of early erythroid progenitors. Following proerythroblast formation, m^6^A-MTase activity becomes dispensable for continued maturation and survival. These findings establish m^6^A -dependent translational regulation as a key determinant of early erythropoiesis and suggest that defective m⁶A signaling may contribute to disorders of ribosome biogenesis such as Diamond– Blackfan anemia^12^.

## Methods

### Human CD34+ HSPC cell culture

Enriched G-CSF mobilized human CD34+ cells were obtained from the Fred Hutchinson Cancer Center CCEH Hematopoietic Cell Procurement core. The material was deidentified prior to being provided for research use. CD34+ HSPCs were grown in StemSpan SFEM II (Stemcell Technologies) supplemented with the following growth factor cocktail (SCF 100 ng/ml, FLT-3 ligand 100ng/ml, TPO 100 ng/ml, IL-6 100 ng/ml) for up to 3 days. EPO was purchased from PeproTech and all other cytokines were purchased from Irvine FujiFilm/Shenandoah Biotechnology.

### In vitro erythroid differentiation

Erythroid differentiation of CD34+ HSPCs was based on the method optimized by Uchida et al.^13^. CD34+ cells were initially cultured for 1-3 days in the media outlined above. The cells were then shifted to erythroid differentiation media consisting of IMDM, 20% Knockout Serum Replacement (Thermo Fisher) 3U/mL EPO, 10ng/mL SCF, 1.0 ng/ml IL-3, 1 uM dexamethasone, and 1 uM estradiol for 5 days. Media was exchanged on day 3 of erythroid differentiation. On day 5 the cells were transferred to erythroid maturation media consisting of IMDM, 20% Knockout Serum Replacement (Thermo Fisher) 3U/mL EPO, 10ng/mL insulin, 0.5mg/ml holo-transferrin, and 2% BSA and cultured until the endpoint of the experiment with media exchanged every 2-3 days. Cell density was maintained between 300,000 and 1.25 million cells/mL.

### RNP nucleofection

Generation of all gene knockouts in CD34+ HSPCs was generated by CRISPR/Cas9 as previously published^14^. (http://www.protocols.io/view/cd34-cell-rnp-nucleofection-q26g7y541gwz/v2) A pair of sgRNAs (Synthego) listed in **Supplementary Table 4** were complexed with sNLS-SpCas9-sNLS Nuclease (Aldevron) in a 2:1 (50 pmol each sgRNA to 50 pmol Cas9) ratio in P3 nucleofector solution (Lonza) for 15 – 20 mins at RT. The RNPs were mixed with 250-400 thousand cells per 96-well sized reaction and nucleofected with program DS-150. Following nucleofection the cells were allowed to recover for a minimum of 4 hours in StemSpan II media (Stem Cell Technologies) with 100ng/ml of TPO, SCF, IL-6 and FLT-3 (Fujifilm Irvine Scientific).

To quantify editing a subset of cells were cultured in the recovery media for 3 days post-nucleofection and genomic DNA was isolated from 50-100 thousand cells with the Quick-DNA MicroPrep Kit (Zymo Research). The isolated gDNA was quantified by nanodrop (Thermo Fisher) and editing was quantified by PCR amplification of regions spanning the two sgRNA cut sites in the edited cells and an unedited control product generated from donor matched unedited cells (Phusion DNA polymerase or Q5 (NEB), or Herculase II (Agilent)). The PCR products were cleaned up with the Monarch Spin PCR & DNA Cleanup Kit (NEB) and sequenced by Sanger sequencing. The Sanger sequencing traces were then run through ICE analysis (Synthego) to obtain an estimate for the indel rate.

### Flow cytometry

HEL cells were stained with APC-CD235a (BD Pharmingen 551336). To monitor CD34+ progenitor cell differentiation status the cells were stained with PE-CF594-CD34 (BD Pharmingen 562383), APC-R700-CD38 (BD Pharmingen 564979), PE-CD123 (BD Pharmingen 554529), Pacific Blue CD45RA (Invitrogen MHCD45RA28), APC-H7-CD41 (BD Pharmingen 561422) and BV786-CD71 (BD Pharmingen 563768). To monitor lineage status cells were stained with APC-CD235a (BD Pharmingen 551336) and PECD71(BD Pharmingen 555537) for erythropoiesis, PE-CD14 (BD Pharmingen 555398) for myeloid cells, and PE-CD41 (BD Pharmingen 557297) and APC-CD61 (BD Pharmingen 564174) for megakaryopoiesis. Cells were analyzed on an BD LSRII or BD Symphony. (Include viability analysis (PI/Annexin V)

### ClickIT Assays

The OPP, EU and EdU assays using Click-iT or Click-iT Plus reagents (Life Tech., C10458, C10340, C10424) were largely done following the manufactures protocol with the following modifications. Erythroid progenitor cells on the indicated day of the assay were incubated with either 10uM OPP or 1mM EU for 1 hour or 10uM EdU for 2 hours. The cells were then harvested and stained with CD235a-PE (BD555570) and CD71-BV786 (BD563768) as described above for flow cytometry. Following surface staining, the cells were fixed and permeabilized using the BD Cytofix/Cytoperm Fixation/Permeabilization Solution Kit (BD554714) and Click-iT labeling was done following the manufactures protocol.

### CFU assays

10^4^ CD34+ HSPCs were plated 4 hours after nucleofection with METTL3 RNPs for lineage specific colony assays (triplicate) in methylcellulose (MethoCult H4230, Stem Cell Technologies) supplemented with 10ng/ml of IL-3, IL-6, G-CSF, GM-CSF, and SCF, and 3U/ml EPO. Colonies were scored after 14 days in culture.

### RNA-seq and splicing analysis

Cells were lysed with for Trizol (ThermoFisher). Total RNA was isolated with the Direct-zol RNA kit (Zymo Research) and quality validated on the Agilent 4200 TapeStation. Illumina sequencing libraries were generated with the NEBNext Ultra II Directional RNA library Prep kit (New England Biolabs, Inc.). Library size distribution was validated using an Agilent 4200 TapeStation (Agilent Technologies). Additional library QC, blending of pooled indexed libraries, and cluster optimization was performed using Life Technologies’ Invitrogen Qubit® 2.0 Fluorometer (Life Technologies-Invitrogen). RNA-seq reads were aligned to the UCSC hg19 assembly using Tophat2^15^ and counted for gene associations against the UCSC genes database with HTSeq^16^. The normalized count data was used for subsequent Principal component analysis and Multidimensional scaling (MDS) in R. Differential Expression analysis was performed using R/Bioconductor package DESeq2^17^. Heatmaps were made using R/Bioconductor package pheatmap (https://CRAN.R-project.org/package=pheatmap).

The rMATS^18^ (http://rnaseq-mats.sourceforge.net/) pipeline version 4.0.2 was used to detect differential alternative splicing events from RNA-Seq data using the parameters -t paired --readLength 50 --cstat 0.0001. The Miso pipeline^19^ was used to analyze the RNA-seq data for alternatively spliced transcripts. First, the expression levels (psi values) were computed for each of the paired end RNA-seq samples individually using ‘miso --run’, followed by calculating the Psi values for each sample using ‘summarise-miso’. Lastly, all pairwise comparisons between the *sgNTC* and *sgWTAP* samples were run using ‘compare_miso’ and the events were fitered using criteria ‘--num-inc 1 --num-exc 1 --num-sum-inc-exc 10 --delta-psi 0.10 --bayes-factor 10’. Only those alternative splicing events that were present in all pairwise comparisons were included in our final results.

### m^6^A-SAC-seq

Total RNA was isolated and either poly(A)-selected or rRNA-depleted. Low-input libraries were prepared from 2–30 ng RNA, following the published m^6^A-SAC-seq protocol. Recombinant MjDim1 was expressed and purified, and the allylic S-adenosylL-methionine (allyl-SAM) cofactor was synthesized as described. m^6^A residues were selectively allyl-labeled by MjDim1 in the presence of allyl-SAM to generate a^6m^6A, then treated with iodine (I₂) to induce intramolecular cyclization of a^6m^6A into an RTdetectable adduct that yields characteristic A→T/C misincorporations during reverse transcription. Libraries were constructed by ligation of sequencing adapters, reverse-transcribed with recombinant HIV-1 reverse transcriptase, and PCR-amplified for Illumina sequencing.

For absolute quantification, synthetic spike-in RNAs bearing defined m^6A stoichiometries were added prior to labeling and carried through library prep to generate per-motif calibration curves converting local mutation rates to site stoichiometry. Reads were adapter-trimmed (Cutadapt), aligned to the reference transcriptome/genome (Bowtie), and processed with the authors’ m^6^A-SAC-seq pipeline to call sites using the expected mutation signature and DRACH context, followed by QC metrics and replication concordance. The method detects m^6^A at single-nucleotide resolution across diverse RNA inputs and tissues with minimal input requirements.

### m^6^A Dot blots

The m^6^A dot blot assay was conducted following our previously established protocol^20^. Briefly, total RNAs were extracted at the indicated time points with miRNeasy Mini Kit (QIAGEN, Cat. #217004). The RNA samples were denatured at 65 °C for 5 minutes in 3 sample volumes of RNA incubation buffer. An equal volume of chilled 20 × SSC buffer (Sigma-Aldrich, Cat. #S6639) was then added, and the samples were spotted onto the Amersham Hybond-N+ membrane (Cytiva, Cat. #RPN303B) using a Bio-Dot Apparatus (Bio-Rad, Cat. # 1706545). After UV crosslinking, the membrane was stained with 0.02% methylene blue (MB), then washed with 1 × PBST buffer, blocked with 5% non-fat milk, and incubated overnight at 4 °C with anti-m^6^A antibody (1:2000, Synaptic Systems; Cat #: 202 003). The membrane was washed with 1 × PBST buffer three times. Subsequently, HRP-conjugated goat anti-rabbit IgG (1:2000; Santa Cruz Biotechnology, Cat #: sc-2005) was applied for 1 hour at room temperature, and the membrane was developed with Amersham ECL Prime Western Blotting Detection Reagent (Cytiva; Cat #: RPN2236).

### m^6^A fluorometric quantification

The manufactures instructions for the m^6^A RNA Methylation Quantification Kit (Abcam) were followed. Human G-CSF mobilized CD34+ HSPCs were in vitro differentiated following the method outlined above either with no treatment, DMSO or 10uM STM2457. At the indicated timepoints RNA was isolated from 250,000 cells as described above for RNAseq and 200 ng of total RNA used per assay. Samples were assayed in triplicate.

### Polysome profiling and analysis

The profiling methodology was based largely on protocols described by Liang and Bellato and colleagues^21,22^. DMSO or STM2457 day 6 in vitro differentiated erythroid progenitors were lysed in hypotonic lysis buffer (10 mM Tris-HCl, pH 8.0, 140 mM NaCl, 1.5 mM MgCl_2_, 0.25% NP-40, 0.1% Trition X-100, 640 U/ml SUPERase-In RNAse inhibitor (Invitrogen), 150 μg/ml cycloheximide, 20 mM DTT) and the cytosolic extract was loaded onto the sucrose gradient. Fractions were collected with a BioComp Gradient Master. The fractions were mixed 1:1 with Trizol LS (ThermoFisher), RNA was isolated with the Direct-zol RNA kit (Zymo Research), and quality validated on the Agilent 4200 TapeStation. Raw sequencing reads were assessed for quality using FastQC (v.0.12.1). Libraries passing quality control were trimmed of sequencing adaptors and aligned to the GRCh38 human genome using STAR2 (v.2.7.3a) and quantified for gene-level expression using HTSeq (v.0.11.1) against the GENCODE v39 gene annotation database to calculate strand-specific read count for each gene. Differential gene expression analysis was performed using DESeq2 (v.1.48.1) to find genes that were transcriptionally regulated using a log_2_fold change of 25% and an adjusted p-value of 0.05. Xtail (v1.2.0) was used to analyze genome-wide translational efficiency by calculating the ratio of polysome-to-monosome for each transcript. Cutoffs were made using a log_2_fold change of 1 ( fold change of 100%) and an adjusted FDR of 0.05 to find transcripts with significant changes at the translational level.

### RT-qPCR

Total RNA was isolated lysing the cells with Trizol and then following the manufactures instructions with the Direct-zol RNA MiniPrep Kit (Zymo Research). The RNA concentration was measured by nanodrop (Thermo Fisher). Oligo dT and SuperScript IV Reverse Transcriptase (Thermo Fisher) were used to generate cDNA from 500ng of RNA and qPCR run with Power SYBR Green Master Mix (Thermo Fisher) on a QuantStudio 7 Flex (Thermo Fisher).

### Gene ontology, Gene set enrichment analysis and Network mapping

Genes sets arising from our genomic data sets were analyzed using GSEA^23,24^, the ToppGene^25^ tool suite, or GeneMANIA network viewer plugin for Cytoscape^26,27^.

## Results

### Early erythropoiesis is dependent on m^6^A-MTase activity

To investigate m^6^A-MTase activity during human erythropoiesis, we utilized a twostep serum-free erythroid differentiation protocol optimized by Uchida et al.^13^ (**Figure 1A**). CD34+ cells were typically expanded for 2-3 days before differentiation and by day 4 of erythroid differentiation, the majority of erythroid progenitors were CFU-E with no CD235+ cells. An average of 42.76±16.28% of CD34+ cells differentiated towards the erythroid lineage (**Supplementary Figure 1A**). On day 4, we transitioned the cells to erythroid maturation media (protocol days 5-17) and by day 6 77.2±7.4% of erythroid cells were expressing CD235. On day 10, 94.0±5.0% of erythroid cells were CD71+/CD235+high (**Supplementary Figure 1A**).

**Figure 1:**
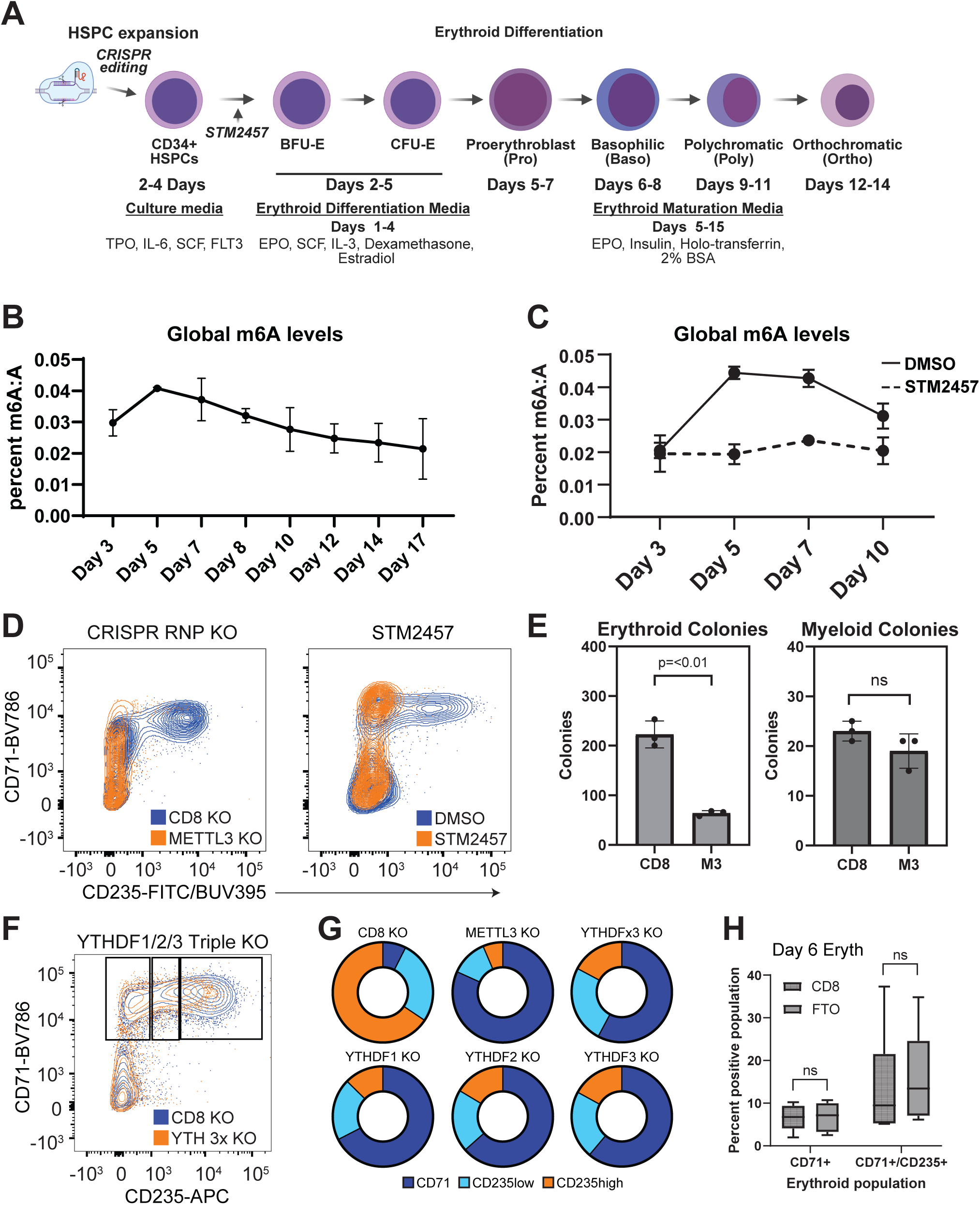
m^6^A methyltransferase activity is required during early erythroid differentiation. (A) Summary of the utilized protocol for in vitro erythroid differentiation of human G-CSF mobilized CD34+ HSPCs. (B) Global m^6^A levels quantified over the course of in vitro erythroid differentiation of human HSPCs by fluorometric quantification (n=3). The day indicates the time relative to the initiation of in vitro erythroid differentiation. (C) Global m^6^A levels following STM2457 treatment quantified over the course of in vitro erythroid differentiation of human HSPCs by fluorometric quantification (n=2). (D) Flow cytometry analysis on day 6 of in vitro erythroid differentiation in control cells or cells with inhibition of m^6^A methyltransferase activity. Inhibition was done by knockout of METTL3 by nucleofection of CRISPR/Cas9 RNPs in CD34+ HSPCs followed by erythroid differentiation or by treating CD34+ cells with 10uM STM2457 in in vitro erythroid differentiation or maturation media. (E) Erythroid and myeloid CFU assay colony counts. Human CD34+ HSPCs were knocked out for CD8 or METTL3 by nucleofection of CRISPR/Cas9 RNPs and colonies counted after 12 days. (F) Representative flow cytometry analysis on day 6 of in vitro erythroid differentiation in CD8 KO control cells or triple KO of YTHDF1,2,3 by nucleofection of CRISPR/Cas9 RNPs in CD34+ HSPCs prior to erythroid differentiation. (G) Quantification of erythroid populations by flow cytometry analysis on day 6 of in vitro erythroid differentiation in control CD8 KO, METTL3 KO cells or cells with single or triple YTHDF gene knockout by nucleofection of CRISPR/Cas9 RNPs in CD34+ HSPCs prior to erythroid differentiation (n=2). (H) Quantification of erythroid populations by flow cytometry analysis on day 6 of in vitro erythroid differentiation in control cells or cells with FTO knockout by nucleofection of CRISPR/Cas9 RNPs in CD34+ HSPCs prior to erythroid differentiation (n=3).

We measured global m^6^A levels over the course of 17 days of in vitro differentiation utilizing a fluorometric assay to quantify m^6^A levels in erythroid progenitors derived from G-mobilized CD34+ cells (**Figure 1B**) and dot blots to quantify levels from CB CD34+ cells (**Supplemental Figure 1B**). In both cases, we saw global m^6^A levels reach their peak as they transitioned from the early BFU-E/CFU-E to proerythroblasts. However, this change in activity did not coincide with similar changes in expression of genes coding for m^6^A methyltransferases, demethylases and reader proteins, although their expression did increase over the course of differentiation (**Supplemental Fig 1C**). The increase in m^6^A levels was due to m^6^A-MTase activity, as treatment with STM2457, a potent m^6^A-MTase inhibitor^28^, kept m^6^A levels at baseline (**Figure 1C**). Furthermore, METTL3 KO^29^ or STM2457 treatment resulted in a near complete block of erythropoiesis during differentiation from the CFU-E to proerythroblast progenitor stages (**Figure 1D**). This was accompanied by a significant reduction in erythroid colony formation, which did not affect myeloid colony formation (**Figure 1E**).

To better understand the regulatory role of m^6^A during early erythropoiesis, we also KO’d m^6^A reader proteins, individually and by family (YTHDF, YTHDC and IGF2BP families), which have been reported to direct different regulatory outcomes for m^6^AmRNA marking (e.g., localization, translation, turnover)^4,30^. The individual and combined KO of the YTHDC and IGFBP families had no effect on erythropoiesis (**Supplemental Fig 1D**). However, individual KO and combined KO of the YTHDF family resulted in a similar block to erythroid differentiation, but with incomplete penetrance compared to m^6^A-MTase treatment (**Figure 2G**). Based on the function of the YTHDF family, this suggests that m^6^A is regulating increased translational and/or RNA degradation. Conversely, knockout of two known m^6^A demethylases, FTO and ALKBH5 had no effect on differentiation (**Figure 1H; Supplmentary Fig 1E**).

**Figure 2:**
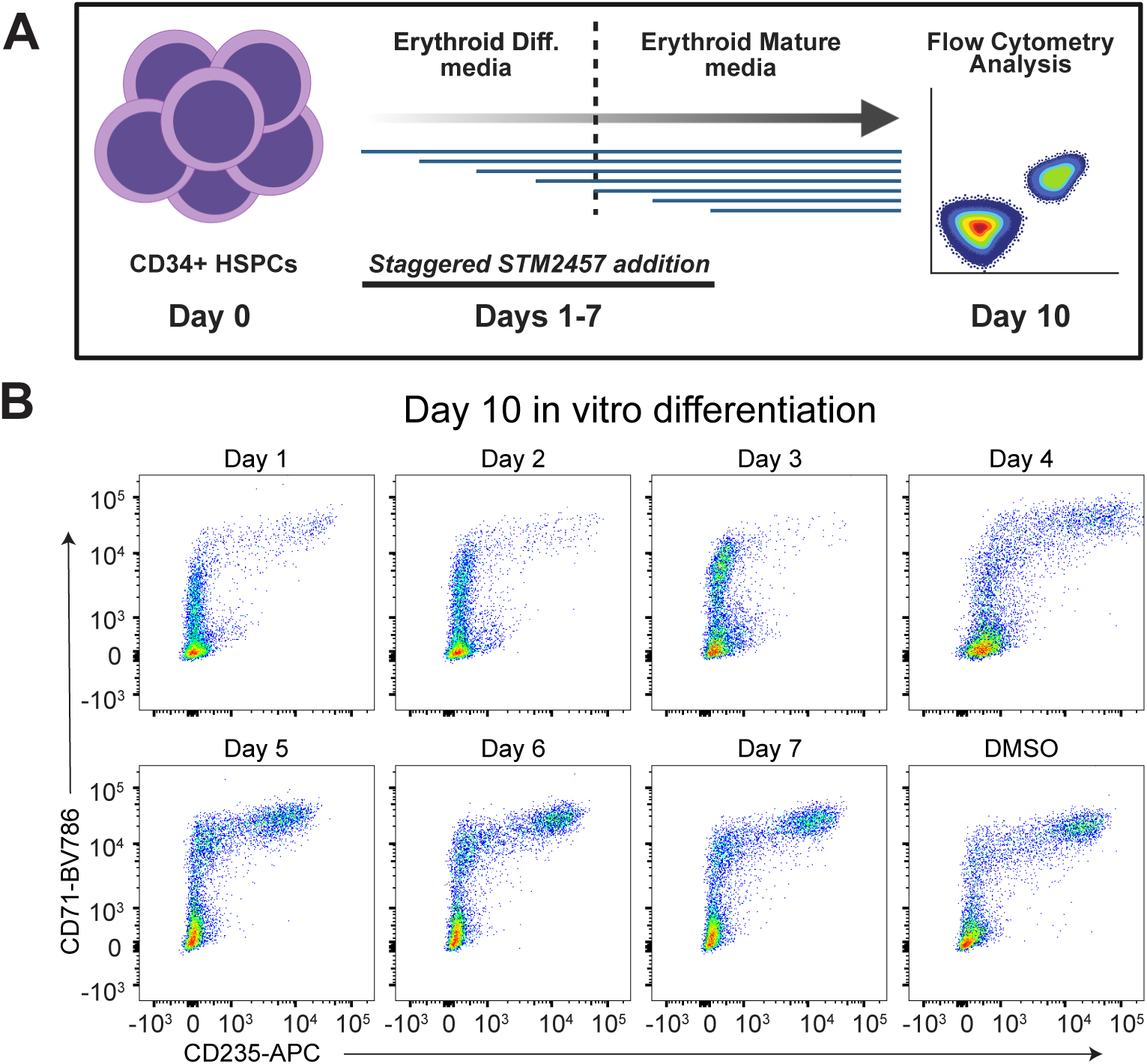
m^6^A methyltransferase activity is only required during an early window for in vitro erythroid differentiation. (A) A diagram of the scheme to investigate staggered STM2457 treatment over the course of in vitro erythroid differentiation. (B) Flow cytometry analysis on day 10 of in vitro erythroid differentiation in control cells or cells with inhibition of m^6^A methyltransferase activity by treating with 10uM STM2457 on the indicated day of in vitro erythroid differentiation.

We next wished to define the timing of the requirement for m^6^A-MTase activity over the course of in vitro differentiation. To this end we staggered STM2457 application over days 1-7 of in vitro differentiation and analyzed erythroid populations on day 10 (**Figure 2A**). The results indicate that m^6^A-MTase inhibition on any of the first three days of erythropoiesis resulted in loss of proerythroblast formation. However, addition of STM2457 on or after day had little or no impact (**Figure 2B**; **Supplemental Fig. 2A**). Thus, a requirement for m^6^A-MTase activity occurs during early erythropoiesis leading up to proerythroblast formation. However, unlike our data from HEL cells, m^6^A mRNA marking is not required to sustain CD235A/GYPA expression or maturation of erythroblasts.

### m^6^A-MTase activity is required for normal proliferation of early erythroid progenitors

To investigate the underlying biological changes in erythroid progenitors leading up to and immediately following the differentiation block, we measured cellular expansion, viability, global DNA replication and cell cycle profiling (**Figure 3A-D**). During the first 3 days of in vitro differentiation, STM2457 treatment did not significantly affect proliferation, but did result in a slowing trend on day 3. A significant plateauing of proliferation occurred on day 4, the final day of the expansion phase of in vitro differentiation (**Figure 3A**). Interestingly, this effect was not accompanied by a significant increase in cell death (**Figure 3B**). However, following transition into the maturation media, there was a trend towards higher cell death in STM2457 treated erythroid progenitors (**Figure 3B**). The data therefore suggest that loss of m^6^A methylation leads to a change in progenitor cell expansion rather than resulting in premature cell death.

**Figure 3:**
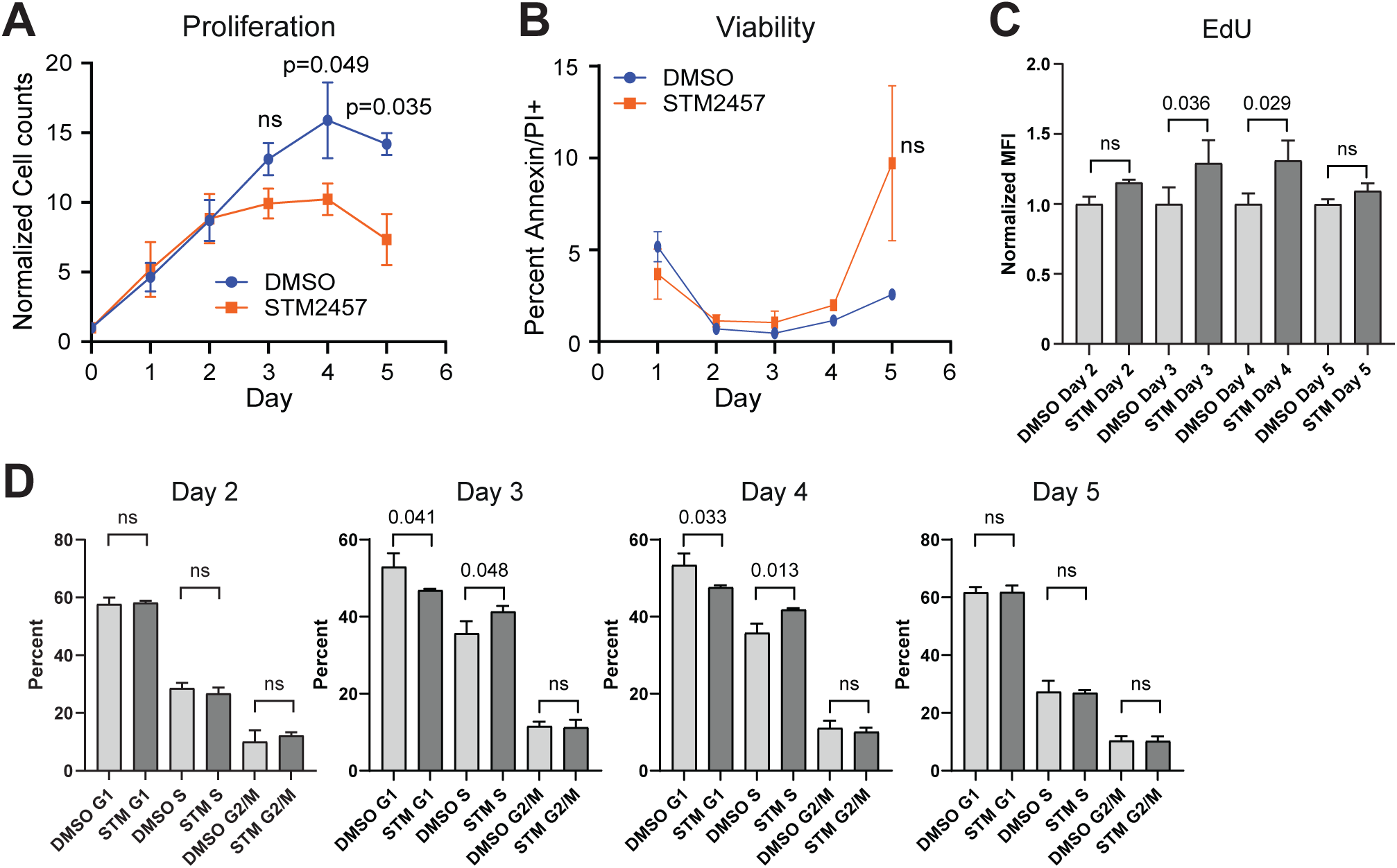
m^6^A methyltransferase activity is required for normal proliferation of erythroid progenitors. (A) Normalized cell counts over the first 5 days of in vitro erythroid differentiation in DMSO or 10uM STM2457 treated CD34+ HSPCs (n=3). (B) Flow cytometry quantification of cell viability by Annexin V and PI staining over the first 5 days of in vitro erythroid differentiation in DMSO or 10uM STM2457 treated CD34+ HSPCs (n=3). (C) Quantification of DNA replication by EdU-ClickIt labeling of all erythroid progenitor cells expressing CD71 on the indicated day of in vitro erythroid differentiation cultured with either DMSO or 10uM STM2457 (n=3)(Day 3 MFI: 2384±38 vs 3086.5±87.5; Day 4 MFI: 1191.5±14.5 vs 1578±88). (D) Quantification of the cell cycle distribution labeling of all erythroid progenitor cells expressing CD71 on the indicated day in vitro erythroid differentiation cultured with either DMSO or 10uM STM2457 (n=3).

We next analyzed the cell cycle profile and DNA replication dynamics of erythroid progenitors from days 2 to 5 of in vitro differentiation. STM2457 treatment caused a clear shift toward increased S-phase entry and DNA synthesis on days 3 and 4, as reflected by both higher EdU incorporation and a corresponding reduction in G_1_-phase cells. Specifically, EdU labeling increased by 30.9 ± 9.9% on day 3 and 31.2 ± 9.8% on day 4 in STM2457-treated cells compared with controls. The proportion of cells in S phase rose from 35.6 ± 2.6% (DMSO) to 41.3 ± 1.2% (STM) on day 3 and from 35.8 ± 2.0% to 41.8 ± 0.3% on day 4, while the G_1_ population decreased from 52.9 ± 2.9% to 46.8 ± 0.3% and from 53.4 ± 2.5% to 47.6 ± 0.4%, respectively (**Figure 3C–D**, **Supplemental Figure 2B**). By day 5, however, both EdU incorporation and cell cycle distribution had normalized to control levels (**Figure 3C–D**, **Supplemental Figure 2B**).

CFU-E undergo multiple rounds of self-renewal cell divisions with the transition to erythroid terminal differentiation (ETD) characterized by progressive shortening of G₁ and a single brief S phase^31^. The CDK inhibitor p57, which regulates S-phase length, plays a causal role in this process^32,33^. Pharmacologic inhibition of S-phase CDKs using roscovitine, an S-phase CDK inhibitor that mimics p57 activity, prolongs S phase and blocks ETD^33,34^. However, combining roscovitine (at 0.5 µM, a dose that rescues expansion of p57 deficient erythroid progenitors without impacting expansion of wild type cells^32^) with STM2457 did not alter the phenotype caused by STM2457 alone (Supplemental Figure 2C-D). This suggests that m^6^A-MTase inhibition either overrides the effects of roscovitine or acts through a distinct mechanism.

### Dynamic regulation of m^6^A mRNA marks during erythropoiesis

To investigate transcript specific changes in m^6^A marking during erythropoiesis, we utilized m^6^A-SAC-seq^35^ to map the m^6^A methylome in human CD34+ HSPCs and in vitro differentiated CD71+/CD235aand CD71+/CD235a+ erythroid progenitors (**Supplemental Fig. 3A, B, C, Supplementary Data 1**). In contrast to antibody-based m^6^A-mRNA capture approaches (e.g., meRIP-seq^36^), m^6^A-SAC-seq provides single base resolution of m^6^A marking and the ratio of m^6^A:A at each site. Consistent with increasing global m^6^A levels during early erythropoiesis (**Figure 4A, Supplementary Data 1**), 29.3% and 16.9% more total m^6^A sites were detected in CD71+ and CD71+/CD235a+ cells respectively compared to CD34+ cells, while average per gene marking increased from 3.0 to 3.8 and 3.9 sites per gene (**Figure 4A**). However, there was only a 1.7% difference in the number of m^6^A marked genes between the CD34+ cells and CD71+ cells, while the CD71+/CD235+ cells averaged 13.3% fewer m^6^A marked genes compared to the other populations (**Figure 4A**).

**Figure 4:**
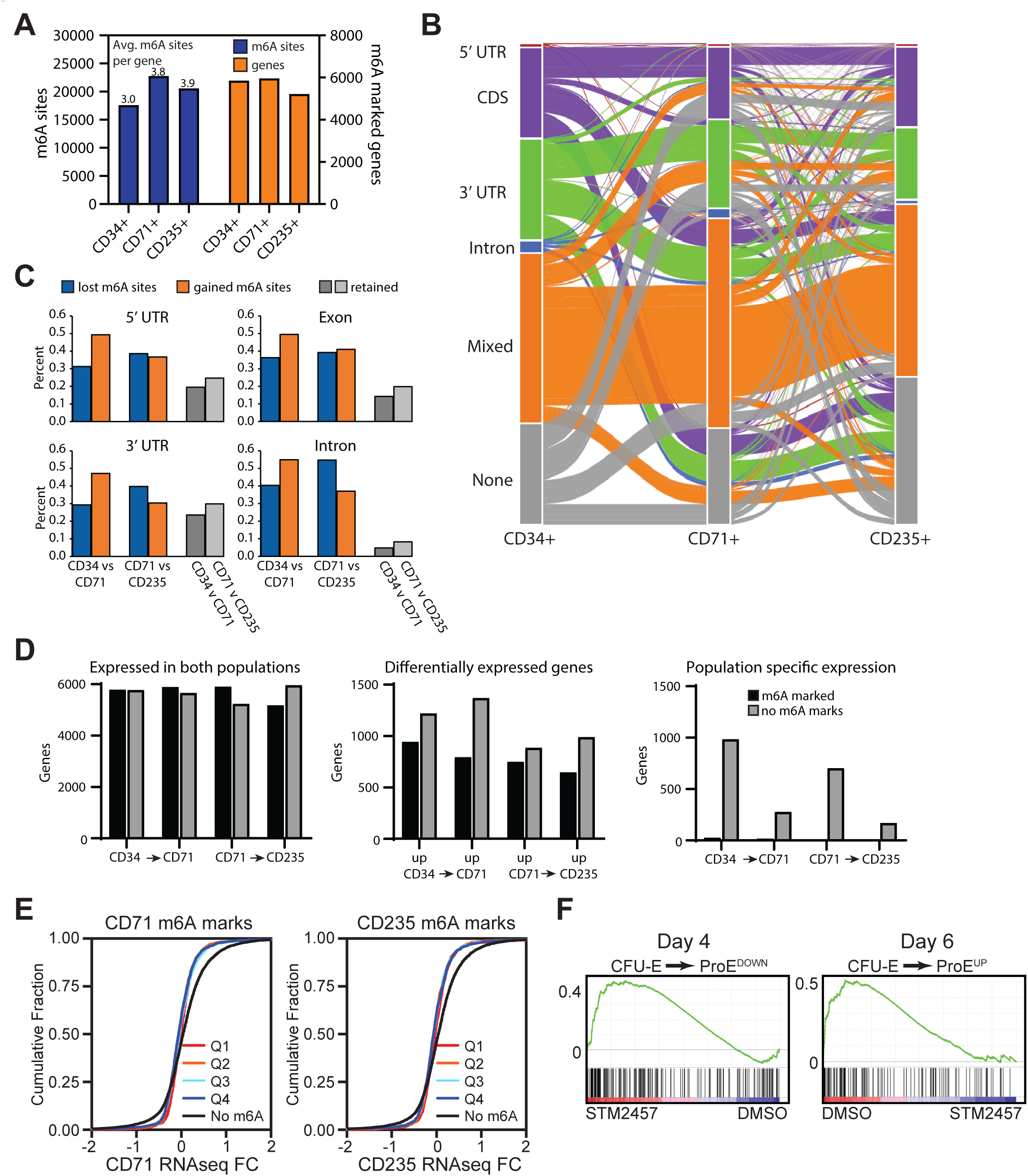
Global and gene specific patterns of m^6^A methylation during in vitro erythropoiesis. (A) Summary of the number of unique m^6^A methylation sites and m^6^A marked genes in HSPCs and erythroid progenitors by m^6^A -SAC-seq (n=3). (B) Alluvial plot showing the dynamics of m^6^A marking at three timepoints of erythropoiesis. Each line represents a single transcript with line color indicating the pattern of m^6^A marking in CD34+ HSPCs. The bars indicate the fraction of transcripts at that stage of erythropoiesis with m^6^A marking in the indicated region of the transcript. (C) The fraction of m^6^A sites in different transcript regions that changed during differentiation between the indicated cell populations. (D) The number of genes with or without m^6^A marking based on changes in gene expression between the indicated progenitor populations. (E) Cumulative distribution plots, separated into quartiles based on the total percentage methylation of all CD71+ or CD235+ erythroid progenitor m^6^A sites within a transcript, for Log2 foldchange in RNA expression comparing DMSO treated and 10uM STM2457 day 6 CD71+ erythroid progenitors. (n=2). (F) GSEA analysis of control and 10uM STM2457 day 4 and day 6 CD71+ erythroid progenitor RNA-seq data using previously described custom gene sets for transcriptional changes during early erythropoiesis^11^.

Examining site specific changes in m^6^A methylation, we detected substantial dynamic regulation of gene specific patterns of methylation (**Figure 4B**). Focusing on genes that are methylated in a minimum of two cell populations, we detected only 50.0-60.4% of genes maintained the same localization of m^6^A methylation across populations. We also detected higher rates of change in gene specific patterns of methylation for genes with only a single mRNA region containing marking versus mixed sites of methylation (3’UTR only: 52.2±3.1%; 5’UTR only: 80.9±7.2%; CDS only: 54.2±3.7% vs mixed: 35.4±7.5%) (**Figure 4B**). The frequency with which m⁶A sites were maintained across differentiation was generally consistent between the CD34⁺ to CD71⁺ and CD71⁺ to CD235⁺ transitions, except for 5′ UTR sites, which were preserved in 26.3 ± 0.7% of transcripts from CD34⁺ to CD71⁺ cells but only 12.0 ± 0.8% from CD71⁺ to CD235⁺ cells (**Figure 4B**).

In CD71+ cells compared to CD34+ cells there were significantly more m^6^A sites gained during progenitor specification than were lost or retained, with 50.2±2.8% of sites gained versus 34.3±4.3% of sites lost and 15.5±7.0% or sites retained (**Figure 4C**). This pattern changed as CD71+ progenitors transition to CD235+ progenitors, with balanced gains and losses in m^6^A sites in the 5’ UTR and exons (38.6% vs 36.7% and 39.2% vs. 41.0%) and a greater loss of m^6^A methylation in the 3’UTR and introns (30.4% vs 39.8% and 37% vs 54.7%)(**Figure 4C**). Since m^6^A methylation exhibits different regulatory roles depending on its position within a transcript, and patterning of m^6^A methylation is associated with different stages of tissue development^37^, the observed changes in m^6^A mRNA patterning during erythropoiesis could reflect changes in m^6^A regulation during lineage commitment versus maturation. Of particular significance for erythropoiesis, genes gaining m^6^A marks going from CD71+ to CD235+ cells are significantly enriched for those involved in hemoglobin synthesis including heme metabolism, mTORC1 signaling and ROS (**Supplemental Fig. 3D**).

### m^6^A-MTase activity does not directly regulate RNA decay during erythropoiesis

Previous studies have shown that m^6^A modification can accelerate mRNA decay during developmental transitions by recruiting RNA-degrading enzymes through YTHDF proteins^4,37,38^. However, when we compared matched RNA-seq and m^6^A -SAC-seq datasets, we found that mRNAs differentially expressed among CD34⁺, CD71⁺, and CD235⁺ populations carried m^6^A marks less frequently than constitutively expressed transcripts (**Figure 4D**). Moreover, transcripts uniquely expressed in each population showed almost no m⁶A methylation (**Figure 4D**). In CD34⁺ cells, these uniquely expressed genes were significantly enriched for pathways related to cell signaling and adhesion (**Supplemental Figure 3E**), whereas no significant gene ontology enrichment was observed for uniquely expressed genes in CD71⁺ or CD235⁺ cells.

To directly assess the impact of m^6^A methylation on gene expression, we performed RNA-seq at two time points: the day before (day 4) and the day after (day 6) the transition to erythroid maturation conditions, with or without STM2457 treatment (**Supplementary Data 2**). At day 4, the numbers of upregulated and down-regulated genes were roughly balanced, whereas by day 6, significantly more genes were transcriptionally upregulated than down-regulated (304 vs. 98) (**Supplementary Data 2**). Notably, and consistent with our prior findings in HEL cells, but contrary to the canonical role of m^6^A in promoting mRNA decay, m^6^A -methylated transcripts were markedly underrepresented among the up-regulated genes vs down-regulated genes (6.25% vs. 21.4%) at day 6 (**Figure 4E**). Gene ontology analysis revealed no pathway enrichment among upregulated genes at day 4, while those upregulated at day 6 were primarily associated with MHC class II function. Overall, we found no correlation between m^6^A methylation levels and transcript abundance (**Figure 4E**), supporting our earlier conclusion from HEL cells that m^6^A marking in this context does not primarily regulate mRNA stability^11^.

However, Gene Set Enrichment Analysis (GSEA) revealed transcriptional changes consistent with the in vitro erythroid differentiation block (**Figure 4F**). Using custom gene sets representing transcriptional programs during early erythropoiesis^11^, we found that STM2457-treated cells retained the CFU-E–like transcriptional program, characterized by persistent expression of genes normally down-regulated during differentiation and reduced induction of genes normally upregulated. Additionally, GSEA identified decreased expression of genes involved in key pathways required for erythroid maturation, including MYC and mTORC1 signaling (**Supplementary Figure 4**).

Together, these data indicate that inhibition of m^6^A methylation results in a transcriptional signature consistent with impaired progression beyond the CFU-E stage. Although we identified a requirement for YTHDF-family readers, known effectors of both mRNA decay and translation, our results suggest that m^6^A marking does not primarily promote mRNA degradation during erythropoiesis. Instead, the YTHDF-dependent effects likely reflect a role for m^6^A in regulating translation.

### m^6^A-MTase activity regulates the translation of genes essential for erythropoiesis

In our previous work using HEL cells, we found that m^6^A methylation promotes the translation of key erythropoietic gene networks, including those encoding ribosomal proteins and the SETD family of histone methyltransferases, chromatin regulators^11^. To determine whether similar translational control mechanisms operate in primary human erythroid progenitors, we performed polysome profiling^21,22^ to assess gene-specific translational changes during in vitro differentiation. Because standard linear-gradient polysome fractionation requires large input cell numbers, we employed a low-input, nonlinear gradient approach as described by Liang et al.^16^. METTL3 knockout was achieved via CRISPR/Cas9 RNP nucleofection, and cells were analyzed after five days of erythroid differentiation (**Supplementary Figure 5A**).

Comparison of the ratio of polysome-associated mRNAs with monosome-associated mRNAs, while excluding those transcriptional changed by RNAseq, revealed 356 translationally down-regulated and 239 translationally upregulated genes in METTL3-deficient cells. Additionally, 61 genes showed concordant transcriptional and translational changes, while 38 displayed opposing trends (**Figure 5A, Supplementary Data 3**). Gene ontology analysis showed no pathway enrichment among translationally upregulated genes, whereas translationally down-regulated transcripts were strongly enriched for ribosomal proteins and mitochondrial energy metabolism/oxidative phosphorylation (OXPHOS) genes, comprising 34 and 59 genes, respectively (**Figure 5B**, **Supplementary Figure 5B**). Consistent with a causal role for m^6^A in translation, translationally down-regulated transcripts were significantly more likely to be m^6^A marked than upregulated ones (48.7% vs. 18.9%, p < 0.0001). The prevalence of m^6^A marking was even higher among ribosomal (64%) and mitochondrial (69%) gene subsets.

**Figure 5:**
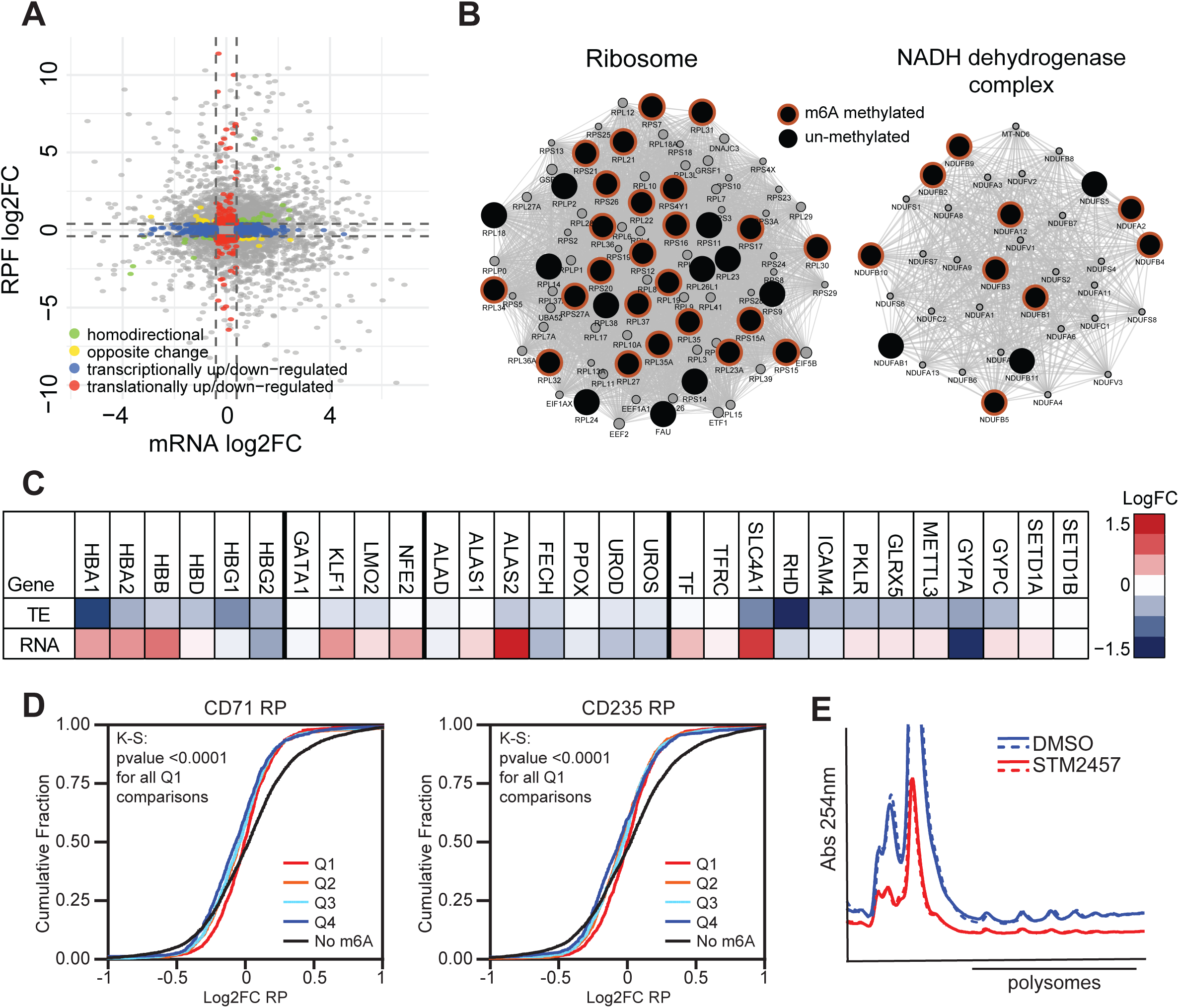
m^6^A-dependent regulation of translation and energy metabolism during erythropoiesis. (A) Translational and transcriptional changes in CD71+ erythroid progenitors on day 6 of in vitro differentiation treated with 10uM STM2457 as measured by polysome fractionation and RNA-seq (n=2). (B) Network maps for interactions enriched in uniquely translationally down genes highlights targeting of ribosomal proteins as well as mitochondrial energy metabolism genes. Large black circles indicate translationally down genes with the orange outline indicating m^6^A marking status in CD71+ erythroid progenitors. (C) A heatmap of the mean translational and transcriptional expression changes of erythroid associated genes in day 6 CD71+ erythroid progenitors treated with 10uM STM2457. (D) Cumulative distribution plots, separated into quartiles based on the total percentage methylation of all m^6^A sites within a transcript, for Log2 foldchange in translation comparing DMSO treated and 10uM STM2457 day 6 CD71+ erythroid progenitors. A leftward-shift indicates reduced translation (n=2, KolmogorovSmirnov Q1 vs. Q2: P value: <0.0001; Q3: P value<0.0001; Q4: P value:<0.0001; None: P value:<0.0001). (E) Representative polysome profiling RNA plot from equal numbers of CD71+ erythroid progenitor cells with and without STM treatment following 5 days of in vitro erythroid differentiation. (n=2)

Several erythroid-associated genes, including HBA1, HBA2, and HBD, as well as erythroid surface proteins, were also translationally down-regulated (**Figure 5C**). Unlike in HEL cells, where m^6^A loss reduced translation of SETD1A/B and disrupted KLF1dependent transcriptional programs^11^, SETD1A/B translation was unaffected here, suggesting that the erythroid phenotype in primary progenitors arises primarily from impaired translation rather than epigenetic dysregulation (**Figure 5C**). Indeed, transcriptional and translational changes were largely decoupled, with fewer than 10% of differentially expressed genes showing concordant directionality (**Figure 5C**).

To explore whether the extent of m^6^A marking correlated with translational regulation, we stratified transcripts by total m^6^A methylation (summed site percentages) and compared translation across quartiles. The most heavily methylated transcripts (top quartile) were significantly less prone to translational down-regulation following STM2457 treatment than other methylated transcripts (**Figure 5D).** The lowest methylation quartile contained 3.6–5.6-fold more translational down-regulation than upregulation genes across CD71⁺ and CD235⁺ progenitors (**Supplementary Figure 5D**). Although the small number of highly methylated and translationally altered transcripts precludes detailed categorization of m^6^A marking patterns associated with translational regulation, this data supports a model in which m^6^A selectively enhances translation during erythropoiesis.

Because METTL3 knockout reduced translation of numerous ribosomal protein genes, we next examined whether this effect resulted in fewer RNA bound ribosome by polysome profiling when inputting equal cell numbers. After five days of differentiation, STM2457-treated CD71⁺ progenitors displayed a 2.2 ± 0.05-fold reduction in monosomes and a 1.6 ± 0.0-fold reduction in polysomes relative to controls (**Figure 5E**), confirming that loss of m^6^A-MTase activity impairs ribosome biogenesis.

### Loss of m^6^A-MTase activity blocks increased translation during early terminal differentiation

Given the reduction in actively translating ribosomes observed upon METTL3 inhibition, we next examined whether global protein synthesis was similarly affected. Using O-propargyl-puromycin (OPP) incorporation assays^39^, we found that day 6 CD71⁺ erythroid progenitors treated with STM2457 displayed a marked decrease in global translation compared with controls (**Figure 6A-C**).

**Figure 6:**
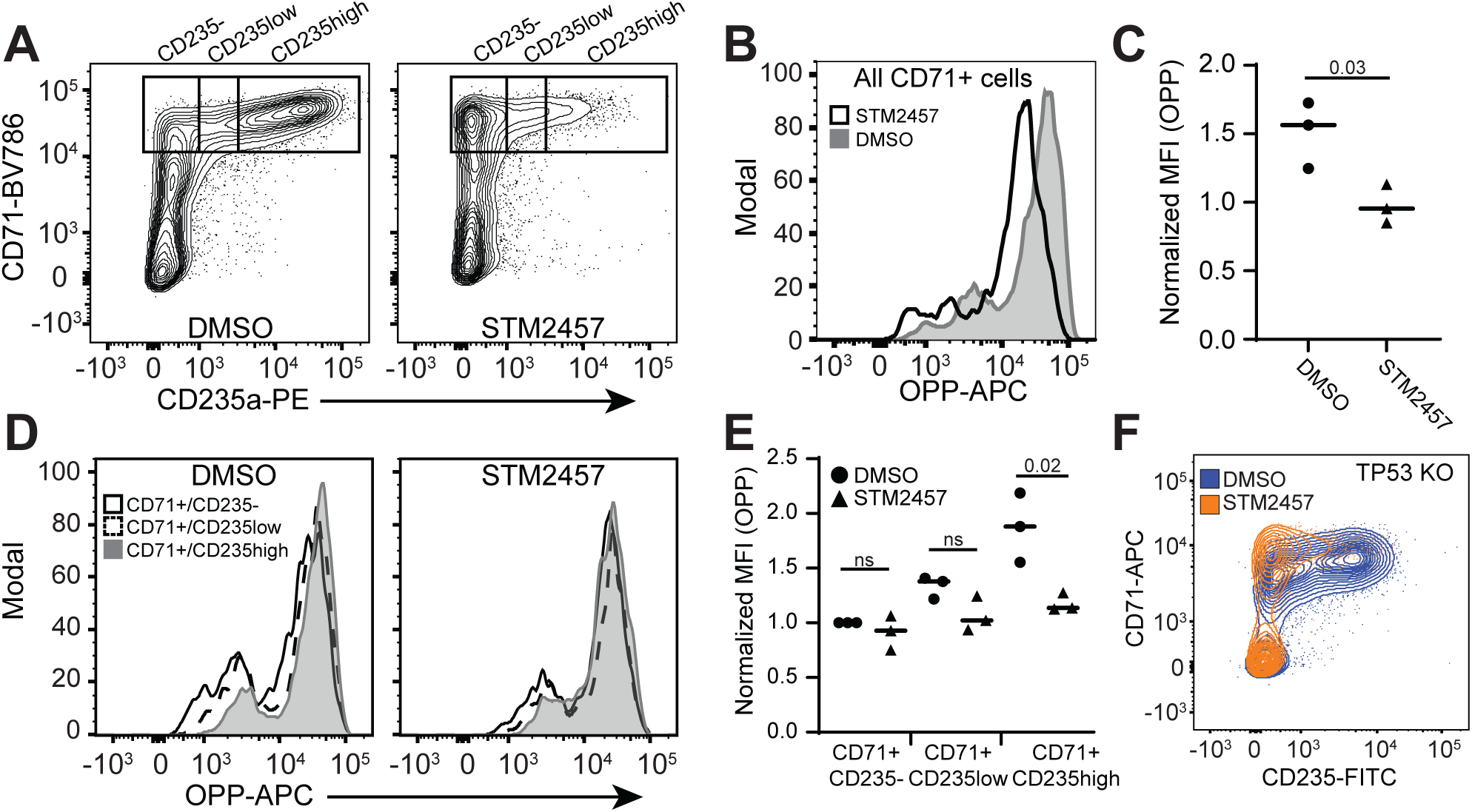
m^6^A regulation of global translation in post-CFU-E progenitors. (A) Representative flow cytometry analysis of erythroid populations on day 6 of in vitro erythroid differentiation in DMSO or 10uM STM2457 treated cells (B) Representative flow cytometry analysis of translation by OPP-ClickIt in all CD71+ erythroid progenitors. (C) Quantification of translation by OPP-ClickIt in all CD71+ erythroid progenitors (n=3). (D) Representative flow cytometry analysis of OPP-ClickIt labeling in the defined erythroid progenitor populations on day 6 of in vitro erythroid differentiation with either DMSO or 10uM STM2457 treatment. (E) Quantification of translation by OPP-ClickIt in the defined erythroid progenitor populations (n=3). (F) Representative flow cytometry analysis on day 6 of in vitro erythroid differentiation culture with either DMSO or 10uM STM2457 treatment and TP53 KO (n=2).

Stratifying erythroid progenitors by CD235a expression revealed a progressive increase in global translation during early terminal differentiation under normal conditions, which was blocked by m^6^A-MTase inhibition (**Figure 6D–E**). These results demonstrate that m^6^A methylation is required to sustain the sharp rise in translational output that accompanies the transition from CFU-E to proerythroblast.

Because loss of m^6^A -MTase activity impairs translation of numerous ribosomal proteins, we asked whether the resulting erythroid defect might be rescued through suppression of the p53 pathway, as previously reported for Diamond–Blackfan anemia (DBA) associated ribosomal deficiencies^40^. To test this, we performed concurrent TP53 knockout using Cas9 RNPs in STM2457-treated cells. However, p53 ablation did not alleviate the erythroid differentiation block (**Figure 6F**), indicating that m^6^A -dependent loss of translation does not trigger a p53-dependent developmental arrest.

Together, these findings establish that m^6^A methylation is essential for the translational up-regulation that supports early terminal erythroid differentiation and that its inhibition phenocopies key features of ribosomopathies such as DBA, albeit in a p53 independent manner.

## Discussion

Hematopoiesis requires precise coordination of transcriptional and translational programs that govern lineage commitment, proliferation, and differentiation. Emerging evidence implicates N^6^-methyladenosine (m^6^A) as a central post-transcriptional regulator of these processes in normal and malignant hematopoiesis. Here, we define a developmentally restricted requirement for m^6^A methyltransferase (m^6^A -MTase) activity during early human erythropoiesis, identifying two distinct erythropoietic windows, during BFU-E expansion and during the CFU-E to proerythroblast transition. Our data reveal that m⁶A levels rise sharply during early erythroid differentiation, peaking in proerythroblasts, and that disruption of METTL3-dependent methylation or YTHDF-mediated m⁶A reading blocks erythroid maturation at the CFU-E stage. These results establish a temporal framework for m^6^A function in human erythropoiesis.

Our findings expand on prior studies demonstrating broad roles for m^6^A in hematopoietic proliferation and lineage specification^5^. In murine hematopoiesis, m^6^A patterning occurs early in hematopoiesis with a static m^6^A landscape playing distinct roles in different progenitor cell populations, whereas our data in human cells indicate dynamic, stage-specific remodeling of the m^6^A methylome during erythroid differentiation^41^. This suggests that m^6^A regulation may be context dependent, governing translation during periods of high biosynthetic demand rather than maintaining fixed epitranscriptomic states. Mechanistically, we show that m^6^A-MTase activity enhances translation of ribosomal, mitochondrial energy metabolism and oxidative phosphorylation (OXPHOS) genes, ensuring maximal global protein synthesis rates and metabolic output coincide with differentiation to proerythroblasts and terminal differentiation^42,43^. Inhibition of m^6^A methylation phenocopies features of ribosomopathies such as Diamond–Blackfan anemia (DBA), including reduced translation, increased erythroid progenitor cell apoptosis, and maintenance of the erythroid expression program in an early progenitor state^12^. We also observe m^6^Aindependent down-regulation of pathways typically suppressed in DBA including heme metabolism and mTOR^12^.

Unlike our prior work in HEL cells, where loss of m^6^A altered translation of epigenetic regulators such as SETD1A/B, in addition to ribosomal proteins, and impaired KLF1-driven transcriptional programs, the effect of m^6^A loss in primary human erythroid progenitors was solely translational. This distinction highlights developmental stage and context-specific roles for m^6^A in hematopoietic cells. The underrepresentation of m^6^A marks on transcripts undergoing expression changes during normal erythropoiesis further suggests that m^6^A acts indirectly to modulate transcriptional programs, likely through its control of translational capacity and cellular metabolism.

In summary, our study establishes m^6^A -dependent translational control as a key mechanism enabling efficient human erythropoiesis. By coupling ribosomal and metabolic gene expression to developmental timing, m^6^A ensures a coordinated increase in biosynthetic capacity during the transition from early erythroid progenitor expansion to terminal erythroid differentiation. These insights not only define a new layer of erythroid gene regulation but also provide a mechanistic link between RNA methylation, ribosome function, and human anemia.

## Supporting information

Suppl Inventory, Figure Legends, and Figures

## Acknowledgements

We thank members of the Paddison, Hsieh, He, Chen, and Torok-Storb labs for helpful advice and comments, Annique Lennon for administrative support, and the Fred Hutch NIDDK-CCEH Cell Processing Core for providing CD34+ HSPCs. Dr. Beverly Torok-Storb, who passed away during manuscript preparation, helped acquire funding for this project and provided critical feedback on experiments. This work was supported by the following NIH grants: NIDDK-P30DK 56465-13, NIDDK-U54DK106829, R01 CA280389 (J.C.), R37 CA230617 (A.C.H), R01 GM135362 (A.C.H.), R01 CA276308 (A.C.H.) and R01 DK119270 (P.J.P.). A.C.H. is also supported by the Larry & Nancy Gordon Endowed Chair for Prostate and Bladder Cancer Research, the Bladder Cancer Advocacy Network, the Nancy & Dick Bernheimer, Matthews Family, Stinchcomb Family, and Thomas & Patricia Wright Memorial Funds. This research was funded in part through the NIH/NCI Cancer Center Support Grant P30 CA015704.

## Authorship contributions

D.A.K. and P.J.P. conceived of the initial idea. D.A.K., J.C., C.H., A.C.H., and P.J.P. designed follow up and mechanistic experiments. D.A.K., R.G., S.L., Y.P. and R.S. performed experiments. C.L.W. and A.W. provided technical assistance. S.A. performed computation analyses and statistical tests. D.A.K. and P.J.P. wrote and revised the manuscript with input from other authors.

