## Supplementary material for "N6-methyladenosine regulation of mRNA translation is essential for early human erythropoiesis": Suppl Inventory, Figure Legends, and Figures

**Supplemental Figure and Table Inventory**

**Supplemental Figures**

**Supplemental Figure 1:** $m^6A$ methyltransferase activity during early erythroid differentiation.

**Supplemental Figure 2:** Cell cycle and DNA replication during early erythropoiesis.

**Supplemental Figure 3:** Mapping $m^6A$ marks during early erythropoiesis.

**Supplemental Figure 4:** GSEA analysis of transcriptional changes in STM2457 treated erythroid progenitors.

**Supplemental Figure 5:** GSEA analysis of transcriptional changes in STM2457 treated erythroid progenitors.

**Supplemental Tables**

**Supplemental Table 1**: $m^6A$-SAC-seq results for CD34+, CD71+ and CD235+ in vitro differentiated erythroid progenitors.

**Supplemental Table 2**: Expression analysis by RNA-seq of DMSO or STM2457 treated erythroid progenitors on days 4 and 6 of in vitro differentiation.

**Supplemental Table 3**: Translational analysis from non-linear polysome fractionation on day 6 of in vitro erythroid differentiation.

**Supplemental Table 4**: sgRNAs and PCR primers used throughout the study.

**Supplemental Figure 1: $m^6A$ methyltransferase activity during early erythroid differentiation.**

(A) Quantification of total erythroid differentiation potential and CD235+ erythroblasts utilizing the differentiation protocol outlined in Figure 1A on days 5 and 10 of differentiation, respectively. (B) Representative flow cytometry analysis of the erythroid populations on the indicated days of in vitro erythroid differentiation of CB CD34+ cells. Dot blot quantification of global $m^6A$ levels on the indicated day with a matching total RNA input blot. (C) qPCR quantification of expression of a select set of $m^6A$ RNA methylation associated gene on the indicated day of in vitro erythroid differentiation. (D) Representative flow cytometry analysis on day 6 of in vitro erythroid differentiation in CD45 or METTL3 KO control cells, or single or triple KO of YTHDC or IGF2BP KO cells by nucleofection of CRISPR/Cas9 RNPs in CD34+ HSPCs prior to erythroid differentiation. (E) Representative flow cytometry analysis on day 6 of in vitro erythroid differentiation in mock edited control cells or KO of ALKBH5 by nucleofection of CRISPR/Cas9 RNPs in CD34+ HSPCs prior to erythroid differentiation.

**Supplemental Figure 2: Cell cycle and DNA replication during early erythropoiesis.**

(A) Quantification of the flow cytometry analysis presented in Figure 2B. (B) Representative flow cytometry analysis of cellular DNA content by DAPI staining and DNA replication by EdU-ClickIt in the indicated erythroid progenitor cell populations on days 2-5 of in vitro erythroid differentiation in DMSO or 10uM STM2457 treated CD34+ HSPCs. (C) Normalized cell counts over the first 5 days of in vitro erythroid differentiation in DMSO or 10uM STM2457 treated CD34+ HSPCs with or without 0.5uM Roscovitine treatment (n=2). (D) Representative flow cytometry analysis on day 6 of in vitro erythroid differentiation culture with either DMSO or 10uM STM2457 and 0.5uM Roscovitine treatment (n=2).

**Supplemental Figure 3: Mapping $m^6A$ marks during early erythropoiesis.**
(A) Representative flow cytometry analysis of erythroid progenitor cell isolation by magnetic beads for generating the erythroid $m^6A$-SAC-seq data. (B) Normalized distribution plots across mRNAs of $m^6A$ marks detected by $m^6A$-SAC-seq in the indicated cell population. (C) Quantification of the genomic feature distributions of $m^6A$ marks mapped by $m^6A$-SAC-seq in the indicated cell population. (D) GO analysis of genes expressed in both the CD71+ and CD235+ populations that gain $m^6A$ sites in the CD235+ cells. (E) GO analysis of genes uniquely expressed in the CD34+ population.

**Supplemental Figure 4: GSEA analysis of transcriptional changes in STM2457 treated erythroid progenitors.** GSEA analysis of control and 10uM STM2457 day 4 and day 6 CD71+ erythroid progenitor RNA-seq data.

**Supplemental Figure 5: GSEA analysis of transcriptional changes in STM2457 treated erythroid progenitors.** (A) Non-linear gradient polysome profiles of day 5 CD71+ erythroid progenitors knocked out for either CD8 or METTL3. (B) GO analysis of translationally down-regulated genes in METTL3 KO CD71+ erythroid progenitors. (C) Venn diagrams comparing transcriptionally and translationally changed genes following

METTL3 KO groups by all genes or genes with $m^6A$ marking in the CD71+ or CD235+ populations. (D) The ratio of genes translationally down to translationally up grouped based on $m^6A$ marking in either the CD71+ or CD235+ population. The numbers above the bars indicate the total number of translationally down genes in the group.

### Supplementary Figure 1

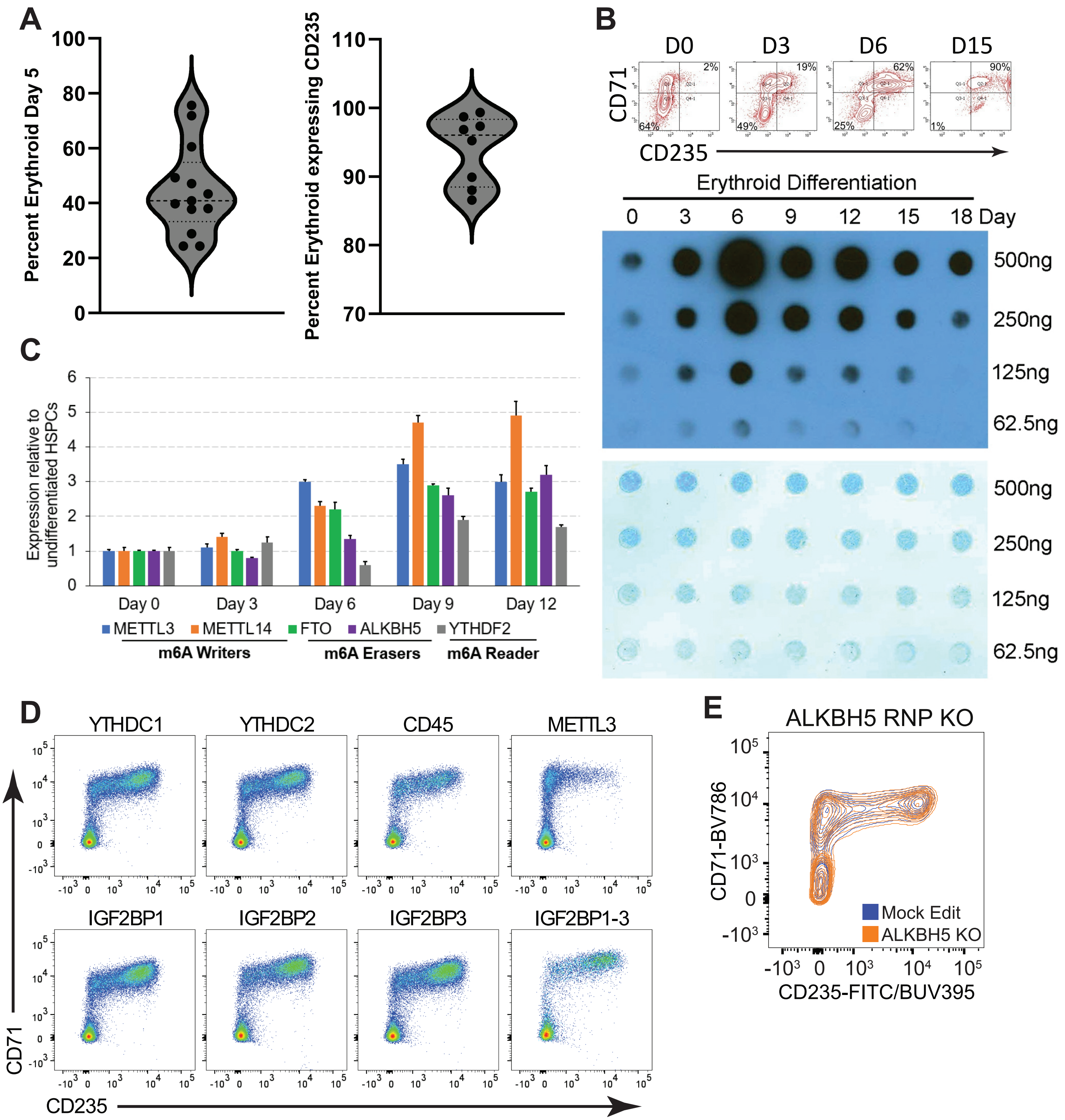

### Supplementary Figure 2

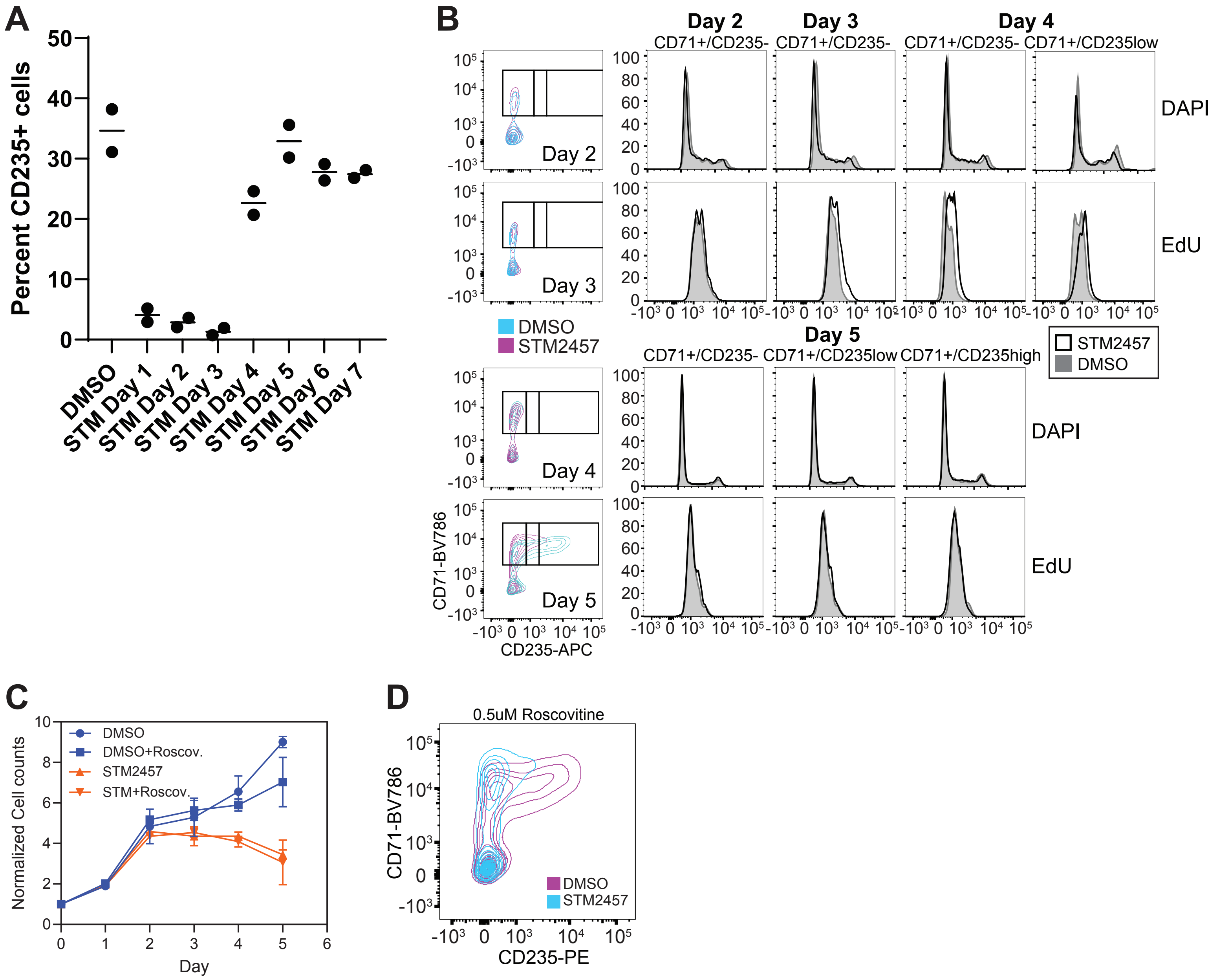

### Supplementary Figure 3

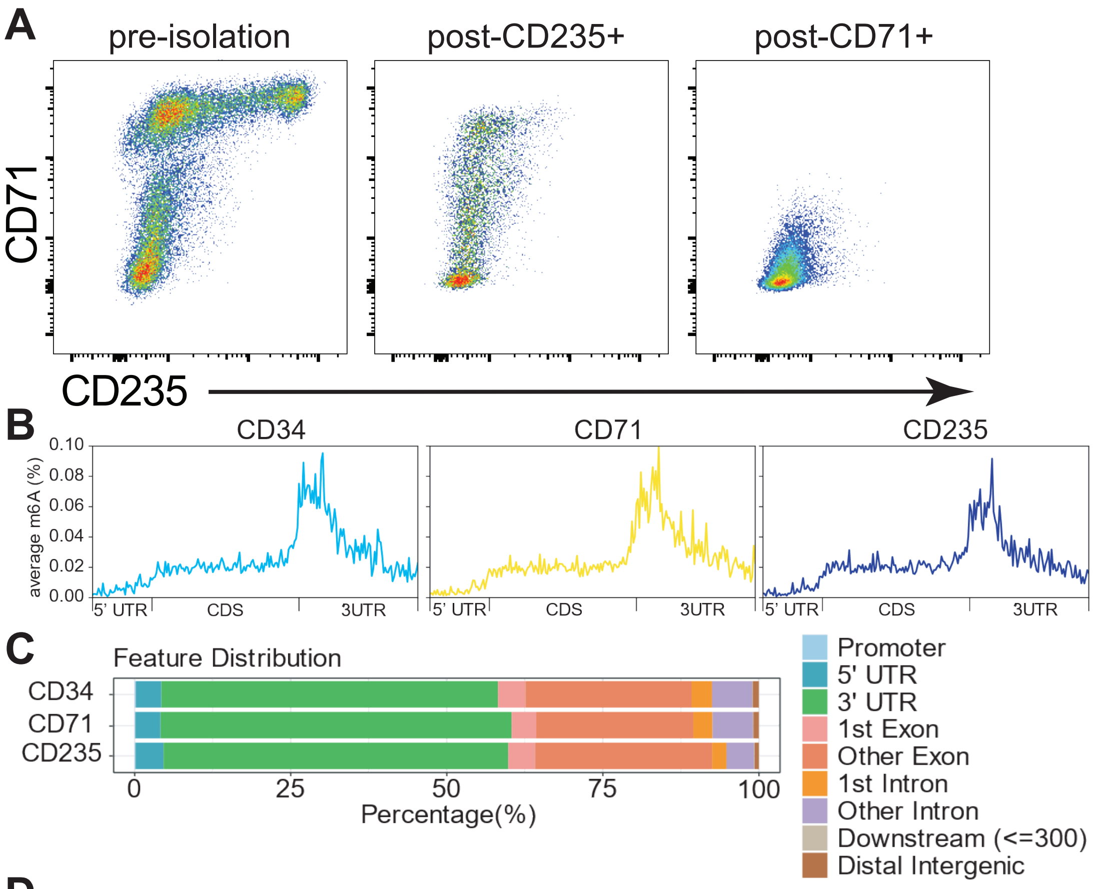


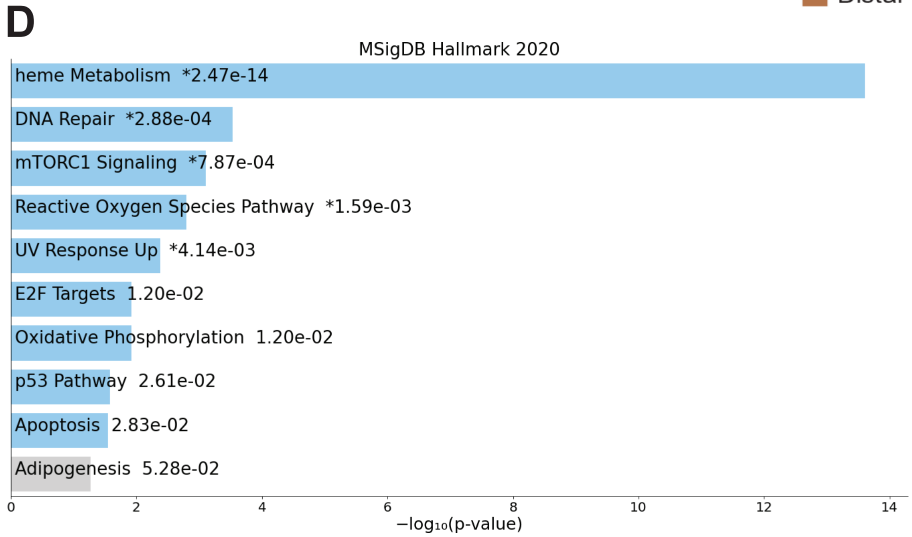


## E

GO: Molecular Function

| ID | Name | pValue | FDR B&H | FDR B&Y | Bonferroni | Genes from Input | Genes in Annotation |
|---|---|---|---|---|---|---|---|
| GO:0005102 | signaling receptor binding | 7.04E-11 | 1.02E-07 | 7.98E-07 | 1.02E-07 | 141 | 1738 |
| GO:0005509 | calcium ion binding | 2.16E-10 | 1.56E-07 | 1.22E-06 | 3.11E-07 | 76 | 749 |
| GO:0005201 | extracellular matrix structural constituent | 6.44E-08 | 3.10E-05 | 2.44E-04 | 9.31E-05 | 28 | 188 |
| GO:0046873 | metal ion transmembrane transporter activity | 3.00E-07 | 1.08E-04 | 8.51E-04 | 4.34E-04 | 48 | 465 |
| GO:0050840 | extracellular matrix binding | 1.24E-06 | 3.58E-04 | 2.81E-03 | 1.79E-03 | 15 | 73 |

GO: Biological Process

| ID | Name | pValue | FDR B&H | FDR B&Y | Bonferroni | Genes from Input | Genes in Annotation |
|---|---|---|---|---|---|---|---|
| GO:0007155 | cell adhesion | 2.49E-17 | 1.76E-13 | 1.66E-12 | 1.76E-13 | 152 | 1675 |
| GO:0007267 | cell-cell signaling | 1.75E-16 | 6.15E-13 | 5.80E-12 | 1.23E-12 | 144 | 1584 |
| GO:0099537 | trans-synaptic signaling | 1.87E-15 | 4.11E-12 | 3.88E-11 | 1.32E-11 | 99 | 939 |
| GO:0098916 | anterograde trans-synaptic signaling | 2.91E-15 | 4.11E-12 | 3.88E-11 | 2.05E-11 | 98 | 931 |
| GO:0007268 | chemical synaptic transmission | 2.91E-15 | 4.11E-12 | 3.88E-11 | 2.05E-11 | 98 | 931 |
| GO:0000902 | cell morphogenesis | 5.62E-15 | 6.59E-12 | 6.22E-11 | 3.96E-11 | 115 | 1194 |

### Supplementary Figure 4

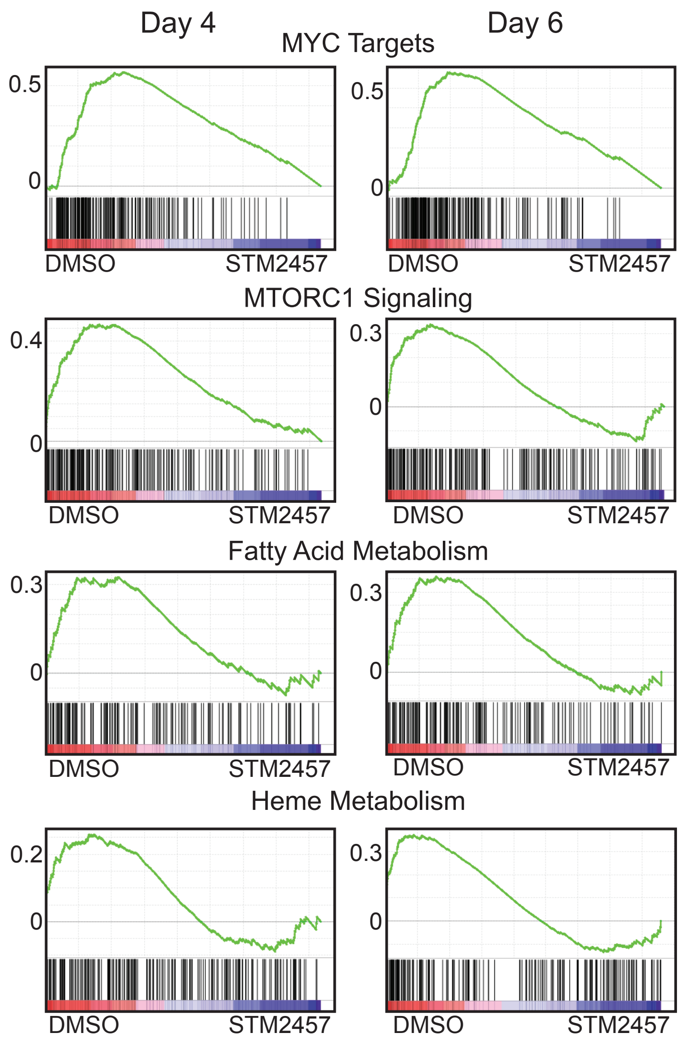

### Supplementary Figure 5

**A**

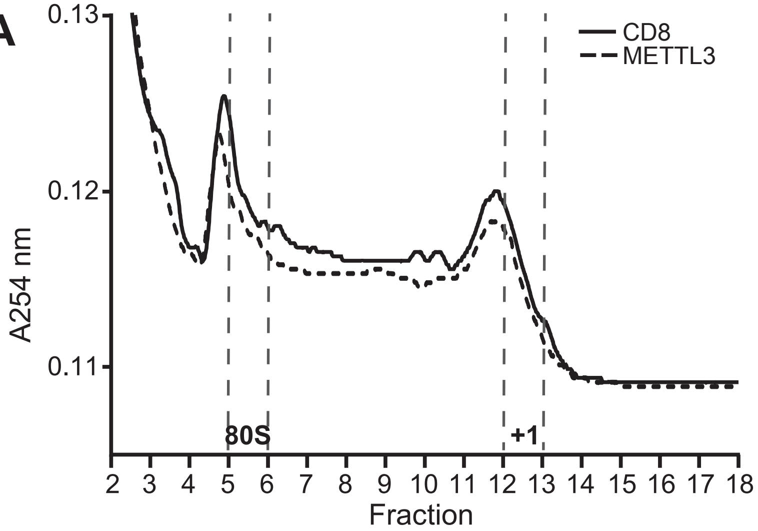


**B**

GO: Molecular Function

| ID | Name | pValue | FDR B&H | FDR B&Y | Bonferroni | Genes from Input | Genes in Annotation |
|---|---|---|---|---|---|---|---|
| GO:0003735 | structural constituent of ribosome | 1.62E-37 | 1.60E-34 | 1.20E-33 | 1.60E-34 | 44 | 181 |
| GO:0005198 | structural molecule activity | 7.18E-16 | 3.55E-13 | 2.66E-12 | 7.10E-13 | 55 | 902 |
| GO:0015453 | oxidoreduction-driven active transmembrane transporter activity | 2.16E-15 | 7.11E-13 | 5.32E-12 | 2.13E-12 | 17 | 68 |
| GO:0009055 | electron transfer activity | 5.83E-14 | 1.44E-11 | 1.08E-10 | 5.77E-11 | 20 | 124 |
| GO:0022853 | active monoatomic ion transmembrane transporter activity | 9.05E-12 | 1.79E-09 | 1.34E-08 | 8.96E-09 | 18 | 126 |
| GO:0008137 | NADH dehydrogenase (ubiquinone) activity | 1.80E-09 | 2.97E-07 | 2.22E-06 | 1.78E-06 | 10 | 41 |

GO: Biological Process

| ID | Name | pValue | FDR B&H | FDR B&Y | Bonferroni | Genes from Input | Genes in Annotation |
|---|---|---|---|---|---|---|---|
| GO:0002181 | cytoplasmic translation | 5.17E-33 | 2.20E-29 | 1.96E-28 | 2.20E-29 | 40 | 181 |
| GO:0140236 | translation at presynapse | 9.91E-25 | 1.79E-21 | 1.60E-20 | 4.21E-21 | 21 | 49 |
| GO:0140241 | translation at synapse | 1.68E-24 | 1.79E-21 | 1.60E-20 | 7.15E-21 | 21 | 50 |
| GO:0140242 | translation at postsynapse | 1.68E-24 | 1.79E-21 | 1.60E-20 | 7.15E-21 | 21 | 50 |
| GO:0006412 | translation | 1.28E-19 | 1.09E-16 | 9.71E-16 | 5.44E-16 | 57 | 826 |
| GO:0042773 | ATP synthesis coupled electron transport | 9.99E-18 | 6.06E-15 | 5.41E-14 | 4.24E-14 | 22 | 109 |

GO: Cellular Component

| ID | Name | pValue | FDR B&H | FDR B&Y | Bonferroni | Genes from Input | Genes in Annotation |
|---|---|---|---|---|---|---|---|
| GO:0044391 | ribosomal subunit | 5.25E-41 | 3.01E-38 | 2.08E-37 | 3.01E-38 | 47 | 193 |
| GO:0022626 | cytosolic ribosome | 1.39E-34 | 3.98E-32 | 2.76E-31 | 7.97E-32 | 36 | 124 |
| GO:0005840 | ribosome | 2.68E-34 | 5.12E-32 | 3.55E-31 | 1.54E-31 | 47 | 264 |
| GO:0005743 | mitochondrial inner membrane | 1.53E-27 | 2.20E-25 | 1.52E-24 | 8.79E-25 | 58 | 599 |
| GO:0015934 | large ribosomal subunit | 6.06E-27 | 6.95E-25 | 4.81E-24 | 3.47E-24 | 30 | 119 |
| GO:0005740 | mitochondrial envelope | 7.08E-23 | 5.80E-21 | 4.02E-20 | 4.06E-20 | 65 | 930 |

**C**

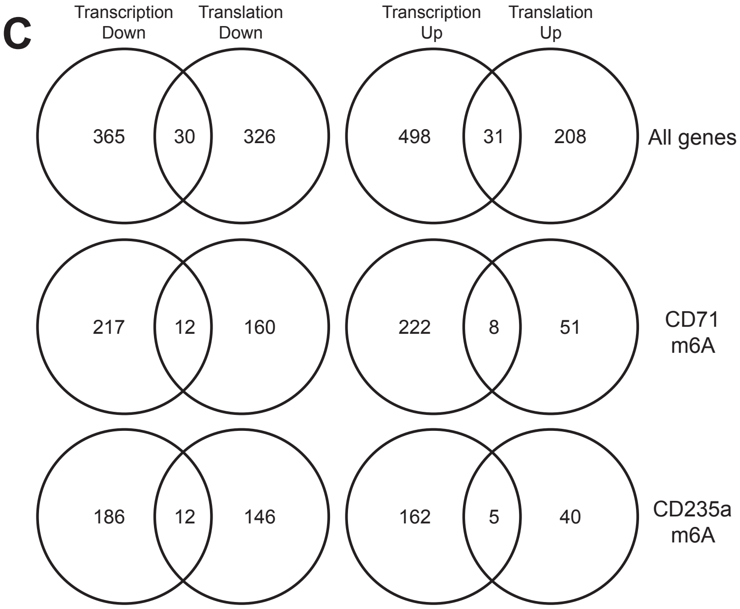


**D**

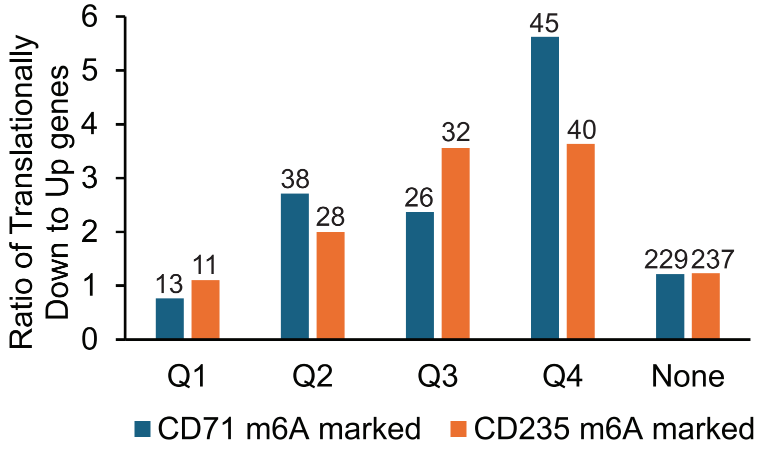
